# Active Microrheology of Intestinal Mucus in the Larval Zebrafish

**DOI:** 10.1101/042994

**Authors:** Michael J. Taormina, Raghuveer Parthasarathy

## Abstract

Mucus is a complex biological fluid that plays a variety of functional roles in many physiological systems. Intestinal mucus in particular serves as a physical barrier to pathogens, a medium for the diffusion of nutrients and metabolites, and an environmental home for colonizing microbes. Its rheological properties have therefore been the subject of many investigations, thus far limited, however, to *in vitro* studies due to the difficulty of measurement in the natural context of the gut. This limitation especially hinders our understanding of how the gut microbiota interact with the intestinal environment, since examination of this calls not only for in vivo measurement techniques, but for techniques that can be applied to model organisms in which the microbial state of the gut can be controlled. We address this challenge by developing a method that combines magnetic microrheology, light sheet fluorescence microscopy, and microgavage of particles, applying this to the larval zebrafish, a model vertebrate. We present measurements of the viscosity of mucus within the intestinal bulb of both germ-free (devoid of intestinal microbes) and conventionally reared larval zebrafish. At the length scale probed (≈ 10*μ*m), we find that mucus behaves as a Newtonian fluid, with no discernable elastic component. Surprisingly, despite known differences in the the number of secretory cells in germ-free zebrafish and their conventional counterparts, the fluid viscosity for these two groups was very similar. Our measurements provide the first *in vivo* measurements of intestinal mucus rheology at micron length scales in living animals, quantifying of an important biomaterial environment and highlighting the utility of active magnetic microrheology for biophysical studies.

## Introduction

The rheological properties of the materials that make up animals are important to physiology (1, 2), development (3), the transport of drugs and biomolecules (4–6), and the activities of commensal and pathogenic microorganisms (7). These properties can depend on the length, time, and shear scales at which they are probed and may furthermore vary in space, mature over time, or respond to biophysical or biochemical stimuli. Although many biological materials can be extracted and characterized *in vitro* via traditional rheological techniques, there are systems for which the complexity of the in vivo environment, the small sizes or volumes of biomaterials present, or the fragility of the substances with respect to changes in their context demand *in vivo* rheological assays. Such is the case with the intestinal fluid of a living animal host, a biomaterial dominated by mucus, a glycopolymer solution or gel whose rheological properties are known to span orders of magnitude depending on, for example, the health of the animal (4).

The rheology of the intestine impacts its vast number of microbial occupants, which must colonize the gut, respond to flows induced by peristaltic motions, and interact with host cells and other microbial cells, all by propelling themselves through the gastrointestinal mucus. In fact, it has been shown *in vitro* that bacteria can alter the rheological properties of mucus directly by chemically modifying it, and it is theorized that this is done specifically to facilitate movement (7). It is also known from studies of germ-free larval zebrafish (*Danio rerio*, shown in Fig. 1 in larval (a) and embryonic (b) stages of development) that the presence or absence of intestinal bacteria modulates the number of mucus producing secretory cells that develop within the host (8). Despite these observations, the rheology of intestinal mucus in germ-free versus conventional animals has never been directly measured.

**Figure 1:**
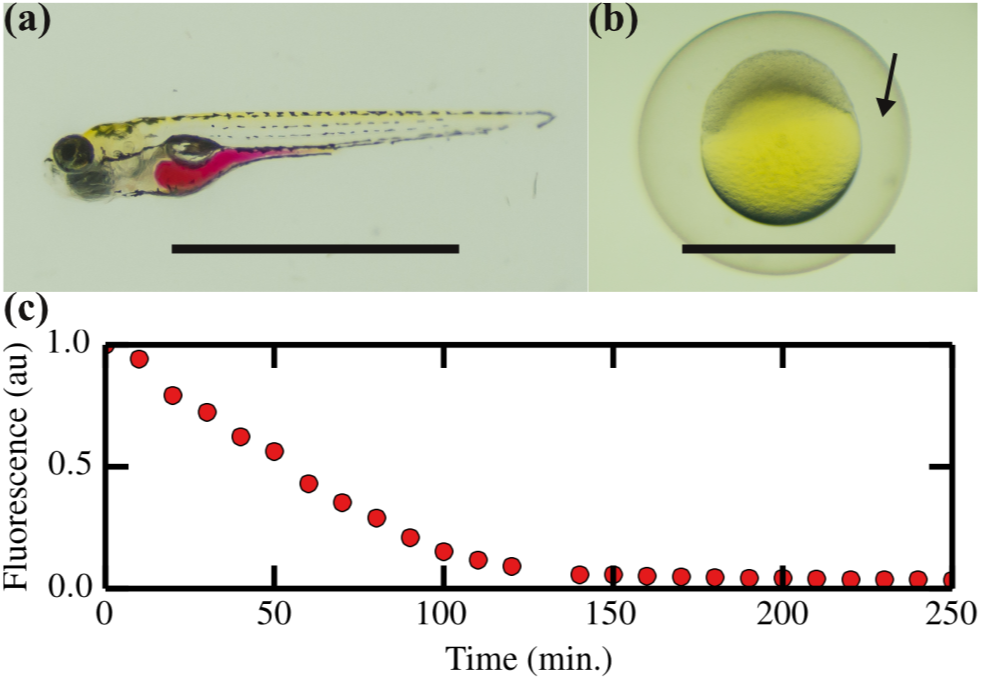
The zebrafish (*Danio rerio*) at two early stages of development and the time scale for external aqueous solutions to exit the intestinal tract. (a) Larval zebrafish at five days post fertilization (dpf). The intestine from bulb to vent is highlighted here by phenol red, which has been orally gavaged. (b) Zebrafish embryo (< 1 dpf) with surrounding chorion. The perivitelline space (indicated by arrow) occupies the region between the embryo and chorion. (c) Fluorescence intensity of a gut gavaged with fluorescent dextran shows that buffer solution is cleared from the gut within two hours post gavage. Scale bars are 1 mm.

Inferring the physical properties of the gastrointestinal environment has been thwarted by a variety of challenges. Especially at early stages in the development of many animals, when the intestine is being colonized by microbes, the volume of the gut is small and difficult to access non-invasively. Moreover, changes in temperature, pH, hydration, and other factors that are unavoidable if mucus is extracted hinder extrapolation from *in vitro* measurements to insights into *in vivo* properties (4, 9). We note also that rheological properties of mucus have been measured *in vitro* to be highly length-scale dependent (4), a consequence of the polymeric nature of mucin glycoproteins, which implies a different effective viscosity at the scales occupied by microbes compared to the macroscopic scales that are typically measured. Passive microrheological techniques that rely on tracking the diffusive displacement of tracer particles have proven useful in a wide variety of studies (10–13). Interpreting tracer response in living organisms, however, can easily be confounded by fluid flows, for example related to circulation or peristalsis. For all these reasons, there exist to date no rheological methods capable of revealing fluid properties in living, developing organisms at the length scales relevant to commensal microbes, despite the importance of quantifying the viscosity of the gut environment. In fact, the only existing method capable of any sort of *in vivo* measurement of gastrointestinal viscosity is based on echo-planar magnetic resonance imaging, which reports viscosity at molecular length-scales and moreover requires highly specialized equipment and large volumes of probe material (“model food”) (14).

To address this, we developed and implemented an approach that combines techniques of microgavage (15), which delivers microparticles specifically to the gut, active microrheology (16–19), which uses external fields to drive particle motions that are easily resolvable over background flows, and light sheet fluorescence microscopy (20–24), which enables rapid, spatially-resolved imaging well inside a living organism. We apply this to measurements of the intestinal rheology of larval zebrafish. Larval zebrafish are a powerful model organism for studying the gut microbiota (8,25–27) due to their biological commonalities with other vertebrates, their optical transparency at young ages, their genetic tractability, and their amenability to gnotobiotic techniques (28), by which larval fish can be kept germ-free (devoid of microbes) or exposed to particular microbial constituents. Using the methods detailed below, we are able to measure the viscosity of the intestinal interior in both germ-free and conventionally colonized zebrafish larvae. These represent the first such measurements to be performed in any living animals. In addition, we measured the viscosity of a different biological fluid: the perivitelline space that separates zebrafish embryos from the chorion, the soft membrane that initially surrounds them. We note that because traditional rheological tools would require sample sizes of 0.10–1.0mL, or the contents of 10^5^–10^6^ larval zebrafish intestines, our microrheological method will prove useful in overcoming similar limitations of sample volume, spatial heterogeneity, and developmental maturation in other systems which remain resistant to standard techniques.

## Materials and Methods

### Theoretical Background

A particle with a magnetic moment m and angular orientation *θ*, located in an external magnetic field of amplitude *B* and direction *β* (as in Fig. 2(b)), will experience a torque of magnitude 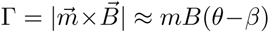, for small values of (*θ* – *β*). This torque tends to align the particle’s magnetic moment with the external field, and the particle must perturb its surrounding environment to do so. For the low Reynolds number system of a colloidal particle in a fluid, a field oscillating as *β*(*t*) = *β*_0_ exp[*iωi*] will result in a particle orientation *θ*(*t*) = *θ*_0_(*ω*) exp[*i*(*ωt* + *ϕ*)]. If the surrounding fluid is characterized by the constitutive law *σ* = *ηγ*̇ for the stress *σ*, viscosity *η*, and strain rate *γ*̇ (i.e., it is a Newtonian fluid), Eq. 1 relates the amplitude of angular displacement of the particle to the driving frequency of the field.

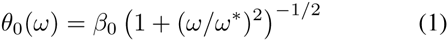

**Figure 2:**
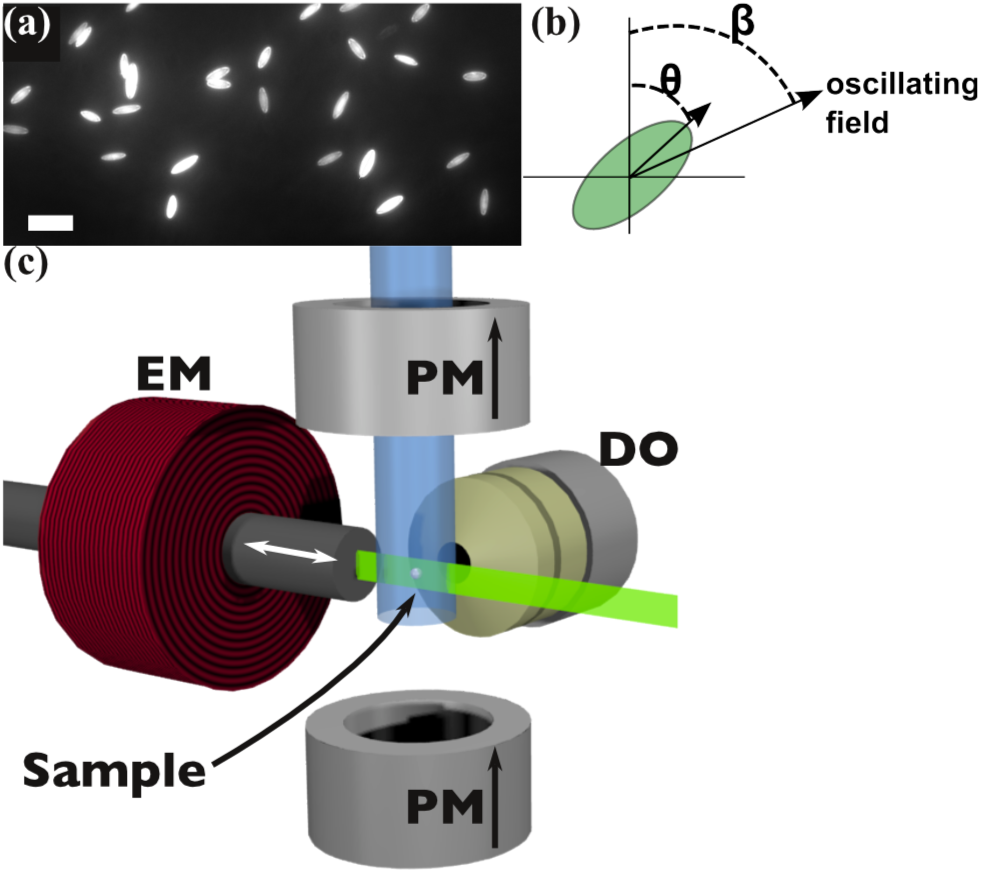
Magnetic probe particles and experimental setup. (a) Fluorescent, super-paramagnetic microspheres are stretched into prolate spheroids (scale bar = 10*μm*). (b) The resulting shape allows visualization of the angular orientation *θ* as the spheroid is driven by an external field with orientation angle *β*. (c) Schematic of the arrangement of magnetic elements and imaging optics, which create the necessary field in the sample volume of a light sheet fluorescence microscope. In this schematic, which excludes the water-filled sample chamber, a vertical field is created by a pair of permanent magnets (PM) while a perpendicularly oriented electromagnet (EM) is used to tilt the direction of the field within the focal plane, imaged through the detection optics (DO). Arrows indicate the direction of magnetic field due to each element, the excitation laser (shown in green) enters from the right, and the sample is held by a capillary that extends through the top magnet into the sample volume.

Here, *ω*^*^ = *mB*/*ξη* and *ξ* is a constant that contains the geometrical factors necessary to make the correspondence between stress/torque and strain/angle (17). Using this expression, the frequency dependence of a particle’s response to an oscillating field can yield a measure of the surrounding fluid’s viscosity. More generally, a material may exhibit non-Newtonian behavior, more fully characterized by the complex shear modulus, as in (17). For our present discussion, we restrict ourselves to Newtonian fluids described by Eq. 1, as this will be shown to sufficiently describe the fluids in question.

### Magnetic Probe Particles

In order to visualize the rotational movement of a particle subject to an oscillating magnetic field, fluorescent, super-paramagnetic polystyrene microspheres (Spherotech, 4.88*μm* diameter) were elongated into prolate spheroids (29, 30). Briefly, particles are first embedded in a thin vinyl film which is fitted onto a mechanical stretcher. This film is then immersed in toluene, which dissolves the polystyrene. Next, the mechanical stretcher is used to elongate the film, deforming the particles along the axis of pull. After removing the toluene, the particles solidify in their new shape and can be recovered by dissolving the vinyl and centrifuging out the particles, which are now shaped as shown in Fig. 2(a, b).

### Microrheology Apparatus and Measurement Procedure

Our experimental setup includes a pair of neodymium magnets located above and below the sample, such that they create a permanent uniform magnetic field directed vertically (Fig. 2(c)). Additionally, a ferrite core electromagnet is mounted to one side of the sample such that, when supplied with a current, the orientation of the field is tilted to an angle *β* with respect to vertical. This is contained within the sample chamber of a light sheet fluorescence microscope with the excitation laser entering from the side opposite the electromagnet, allowing the generation of the magnetic field described above, whose vector orientation oscillates within the imaging focal plane.

In Eq. 1, *m*, *B*, and *ξ* are unknown system parameters, however, the quantity *mB*/*ξ* does not vary appreciably for different particles driven by the same field, as seen in Fig. 3(a). We therefore use a fluid of known viscosity to determine this quantity and calibrate the system for subsequent measurements on fluids of unknown viscosity. For this calibration, we independently measured the viscosity of a water/glycerol mixture by tracking the diffusion of 200 nm diameter, spherical, colloidal particles using an inverted wide-field fluorescence microscope and applied the Stoke–Einstein–Sutherland relation to determine the mixture’s viscosity. Measurements of *θ*_0_(*ω*) for actively driven ellipsoids in the same glycerol solution, fit to Eq. 1, provided the necessary information to calibrate the method.

**Figure 3:**
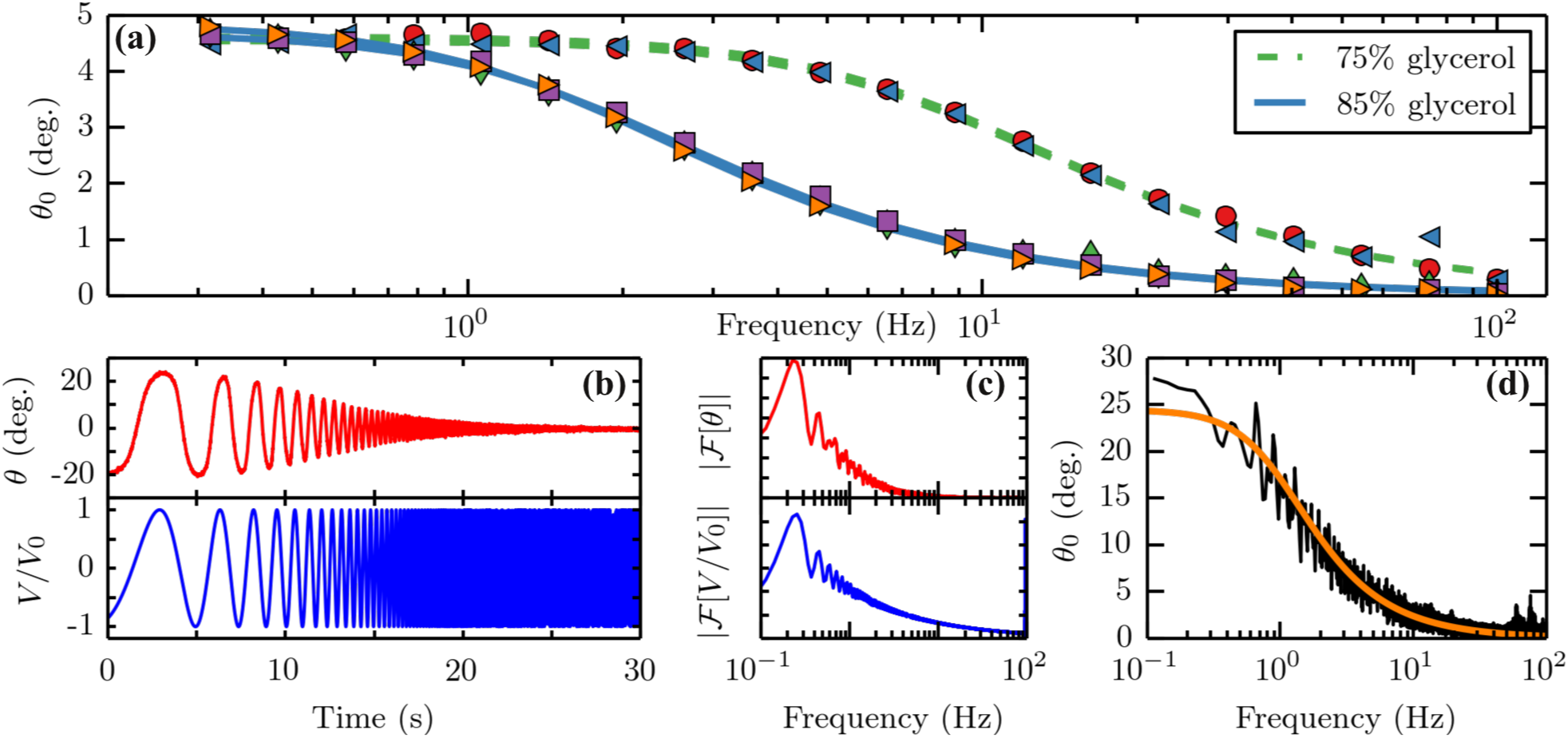
Frequency response in a water/glycerol calibration fluid. (a) *θ*_0_ is measured at discrete frequencies in two different water/glycerol preparations. The curves are fits to Eq. 1. The consistency between different probe particles demonstrates the uniformity in their shape and magnetization, allowing calibration using the known viscosity of these solutions. (b-d) Particles are driven with a continuously increasing frequency, allowing a rapid measurement. First, the orientation of the particle is measured (b, top) in response to the externally supplied electromagnet voltage (b, bottom). Next, Fourier transforms are computed for the two signals (c). Finally, taking the absolute value of the ratio of these computed signals yields *θ*_0_(*ω*) (d), which can be fit to Eq. 1 to obtain a measure of viscosity *η*.

The most obvious procedure to measure *θ*_0_ as a function of diving frequency *ω* would be to incrementally drive the particle through a series of discrete frequencies while recording high speed video of the resulting motion. Once the orientation is extracted from the video, the amplitude and phase of the signal can be computed by using the known driving signal and the orthogonality of sine waves over an integral number of periods (this is the procedure followed in collecting the data in Fig. 3(a)). When performing a measurement in a system such as the intestinal bulb of a larval zebrafish, however, this procedure is too time consuming, as other sources of fluid flow (e.g. peristaltic motion in the gut) which may occur during data collection corrupt the particle motion and invalidate the measurement. We therefore drive the magnetic field with an exponentially chirped signal of the form *ω*/2*π* ≡ *f* = *f*_0_*k*^*t*^ with *k* ≡ (*f*_*f*_/*f*_0_)^1/*T*^ determined by the initial frequency *f*_0_, final frequency *f*_*f*_, and the time (*T*) taken to sweep from the former to the later.

Fluorescence images are captured with a scientific camera (Pco.edge, PCO AG) at ≈ 200 frames per second and the particle orientation is tracked using custom code written using OpenCV (31). A triggering signal from the camera ensures that the driving signal begins its frequency modulation in sync with image acquisition and a time stamp encoded in each image enable us to account for occasionally dropped frames. The spectra of the input and output signals are related by the system transfer function *H*(*ω*), as:

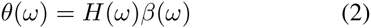

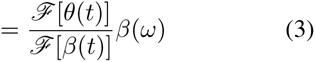

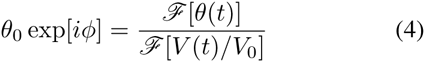

where ℱ denotes the Fourier transform and *V* = *V*_0_ exp[*iωt*] is the voltage applied to the electromagnet, to which *β* is proportional. Using this relationship, *θ*_0_ is calculated from the measured orientation of the particle and the known driving voltage on the electromagnet and can be fitted to Eq. 1 in order to extract *ω*^*^ and, therefore, the viscosity *η*, as summarized in Fig. 3(b-d).

To estimate the uncertainty in *ω*^*^, we simulated *θ*(*t*) signals for various values of *ω*^*^, adding various levels of Gaussian distributed noise. These signals were then used to compute the transfer function from Eq. 4, which was fit to Eq. 1. The inferred value of *ω*^*^ was compared with its actual value, giving a mapping from noise level and *ω*^*^ value to expected error. Data from experiments were assigned uncertainty values based on their individual noise levels and value of *ω*^*^.

### Microinjection and Microgavage of Zebrafish Embryos and Larvae

In order to perform our microrheological measurements, we must first introduce magnetic tracer particles into the system of study. This is easily done for the perivitelline space by piercing the chorion with a micro-pipette and injecting a solution in which particles have been suspended. Getting these particles into the intestinal bulb of zebrafish larvae is more difficult, since we do not wish to cause lethal injury by a similar injection technique. In principle, it could be done by doping a food source with the magnetic particles prior to feeding the zebrafish larvae. Such a strategy, however, would likely change the gut environment substantially by the introduction of food matter. (At these ages, larvae can subsist on yolk material.) Furthermore, we would like to be able to preserve the germ-free or gnotobiotic nature of the specimens in order to address physiological questions in which the properties of mucus may change, and it is at present impossible to prepare germ-free, normally fed larval zebrafish. We therefore adopted a recently developed technique of orally gavaging zebrafish larvae (15), where fluid is delivered to the intestinal bulb by inserting a micro-pipette tip into the esophagus of the fish an injecting a small volume of material. Using this method, we are able to introduce a volume of fluid small enough that it does not visually displace mucus from the gut (as seen by mucus autofluorescence), but carries with it the desired tracer particles. In order to control for the fluid that accompanies the tracer particles, we estimated the time required for the gut environment to return to its pre-injection composition by gavaging zebrafish larvae with an aqueous solution of fluorescent dye and measuring its brightness in the gut. As seen in Fig. 1(c), such a solution is not detectably present in the gut two hours after the initial injection. We therefore restrict our data collection to times that are at least two hours after gavaging tracer particles into the fish. Notably, in many cases, the particles do not remain in the fish for such long periods, and measurement is therefore impossible.

### Animal Use Protocols

The University of Oregon Institutional Animal Care and Use Committee approved all work with vertebrate animals.

Experiments were carried out in either the intestinal bulb of zebrafish larvae (5 days post fertilization, Fig. 1(a)) or the perivitelline space of zebrafish embryos (< 24 hours post fertilization, see Fig. 1(b)).

## Results and Discussion

We gavaged or microinjected ellipsoidal microparticles into the intestinal bulb of conventionally-reared and germ-free zebrafish larvae, as well as the perivitelline space of zebrafish embryos, and performed the active microrheology procedure described in Methods, using the calibration illustrated in Fig. 3. In each of these environments, the response curves are well fit by the form expected of a Newtonian fluid (Eq. 1), as shown in Fig. 4. The length-scale probed by the microparticles is similar to the size of bacteria, implying that bacteria occupying the larval intestinal space will experience a predominantly viscous fluid environment. Any elastic component of the fluid would manifest as a nonzero lower limit to the oscillation amplitude at high frequencies (17), which is not seen in any of the data collected (e.g. Fig. 4). This *in vivo* observation is in contrast to *in vitro* measurements of human cervicovaginal mucus, which show non-Newtonian behavior at length scales above a few hundred nanometers (4), suggesting either that *in vitro* preparation artifacts, developmental stage, or animal species has a major impact on rheology.

**Figure 4:**
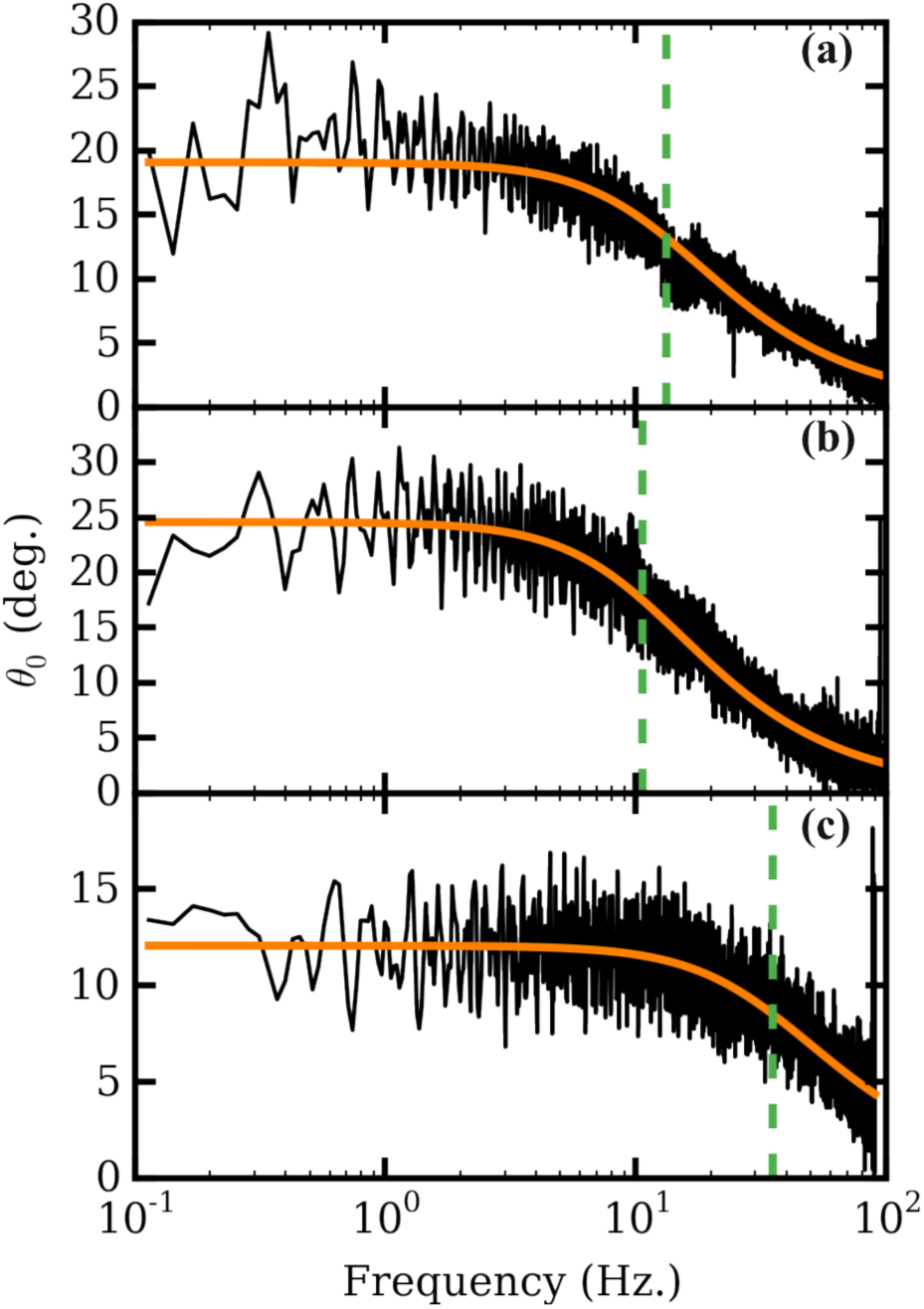
Amplitude response curves. Typical response *θ*_0_(*ω*) of intestinal mucus in conventional (a) and germ-free (b) zebrafish larvae, and the perivitelline fluid of zebrafish embryos (c). In each case, the black line is experimental data, fit to Eq. 1 by the orange curve, with the fit parameter *ω*^*^ indicated by the vertical dashed line.

Repeating this measurement over several specimens, we compute the weighted mean of the viscosity in these three systems to be 4.72 ± 0.51 cP, 5.71 ± 0.95 cP, and 3.12 ± 0.95 cP for the intestinal fluid of conventional and germ-free intestines, and for the perivitelline fluid, respectively (as shown in Fig. 5; N=10, 5, and 12 samples, respectively). Uncertainties are the standard error of the weighted mean, as described in Methods. The viscosity of the perivitelline fluid is about 3 times that of water, providing a quantification of the decades-old observation that the perivitelline space is “filled with a viscous fluid” (32).

**Figure 5:**
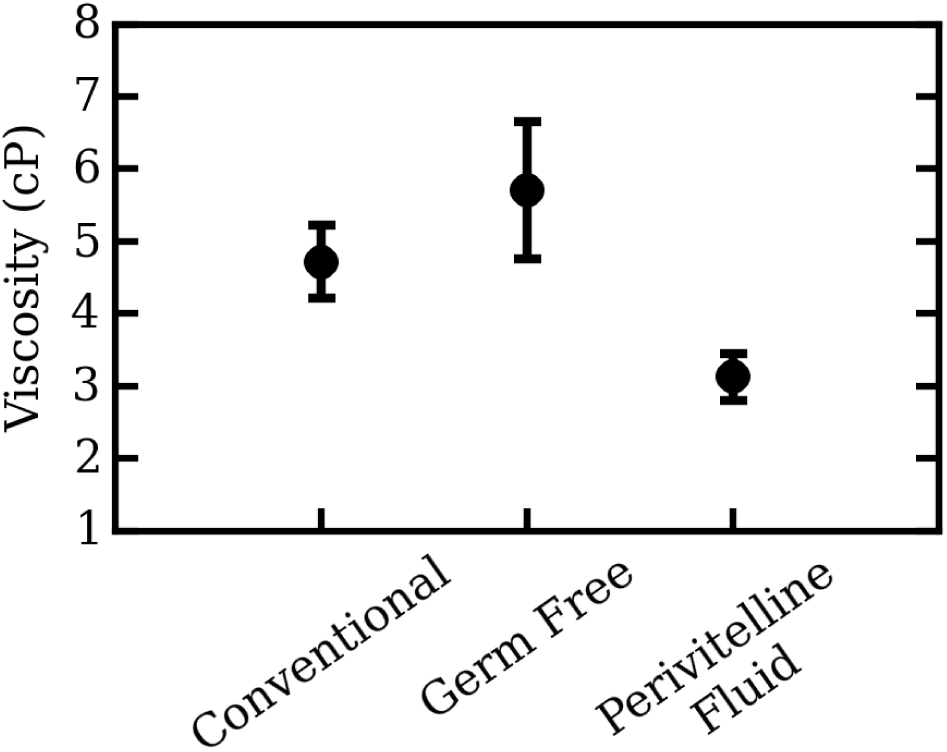
Viscosity measurements. Measured viscosities of intestinal mucus in conventional and germ-free zebrafish larvae, and the fluid occupying the perivitelline space of zebrafish embryos. Circles indicate the weighted mean values, and bars show standard errors of the weighted mean (with calibration error included).

We find values of mucus viscosity that are about 5 times larger than that of water, with a slight enhancement in germ-free relative to conventional fish. The similarity of the values in the two cases is perhaps surprising, as zebrafish which have been derived germ-free posses about 50% fewer mucin-producing secretory cells than conventional fish (8). Our observation may reflect the developmental stage of the animal, which at the time of experiment is still nutritionally sustained by a yolk and has yet to ingest solid food, which would likely stimulate the activity of secretory cells. In either microbial state, our measured viscosities are much lower than those of previously mentioned *in vitro* studies of mucus, which report values at low shear rate on the order of 100 cP (4). Clearly, understanding these differences will require insights into the influence of developmental stage, species, physiological state, and measurement method on mucus rheology.

Mucus provides a rich set of biophysical mysteries that are of both fundamental and practical importance. Our understanding of this fluid is still in its infancy, and is based largely on in *vitro* assays that may poorly mirror *in vivo* environments, and that are prone to artifacts. For example, in contrast to measurements of reconstituted mucus that report a lowering of viscosity by *Helicobacter pylori* (7), other studies report an *increase* in mucus viscosity following *H. pylori* infection, ascribed to possible differences in extraction and handling methods (9).

We suggest that the methodology we describe here offers a path to *in vivo* measurements that will be crucial for making sense of mucus biophysics. This opens the doors to examinations of mucin response in relation to host or microbial physiology, and comparisons to the rheologies of other complex fluids. Notably, this path is so far rather low-throughput and technically challenging, and would be well served by further methodological improvement. More generally, the combination of active microrheology and light sheet fluorescence microscopy should be applicable to a wide variety of small, structurally complex biophysical environments such as circulatory or lymphatic vasculatures, or lumenal spaces of various tissues and organs, that have yet to be quantitatively investigated.

## AUTHOR CONTRIBUTIONS

M.J.T. and R.P. designed the research; M.J.T. performed the research and analyzed data; M.J.T. and R.P. wrote the article.

## ACKNOWLEDGEMENTS

We thank Rose Sockol and UO Zebrafish Facility staff for fish husbandry, and Karen Guillemin, Eric Corwin, Judith Eisen, Kyle Welch, and Ryan Baker for useful discussions and comments. Research reported in this publication was supported by the National Science Foundation under Award no. 0922951, the M. J. Murdock Charitable Trust, the University of Oregon through a faculty research award, and by the National Institutes of Health as follows: by the NIGMS under award number P50GM098911 and by the NICHHD under award P01HD22486, which provided support for the UO Zebrafish Facility. The content is solely the responsibility of the authors and does not represent the official views of the NIH, NSF or other funding agencies.

